# Machine learning methods to reverse engineer dynamic gene regulatory networks governing cell state transitions

**DOI:** 10.1101/264671

**Authors:** P. Tsakanikas, D. Manatakis, E. S. Manolakos

## Abstract

Deciphering the dynamic gene regulatory mechanisms driving cells to make fate decisions remains elusive. We present a novel unsupervised machine learning methodology that can be used to analyze a dataset of heterogeneous single-cell gene expressions profiles, determine the most probable number of states (major cellular phenotypes) represented and extract the corresponding cell sub-populations. Most importantly, for any transition of interest from a source to a destination state, our methodology can zoom in, identify the cells most specific for studying the dynamics of this transition, order them along a trajectory of biological progression in posterior probabilities space, determine the "key-player" genes governing the transition dynamics, partition the trajectory into consecutive phases (transition "micro-states"), and finally reconstruct causal gene regulatory networks for each phase. Application of the end-to-end methodology provides new insights on key-player genes and their dynamic interactions during the important HSC-to-LMPP cell state transition involved in hematopoiesis. Moreover, it allows us to reconstruct a probabilistic representation of the “epigenetic landscape” of transitions and identify correctly the major ones in the hematopoiesis hierarchy of states.

## INTRODUCTION

Stem cell populations exhibit heterogeneous gene expression profiles, indicative of the dynamic changes occurring during differentiation when individual cells transition from one phenotypic state to another^1^. Although the source of this "biological noise” remains unclear, it seems to reflect the variability in cell signaling mechanisms^2^. Traditionally, cell biology studies have drown conclusions on cell behavior based on population statistics^3^. However, cell fate decisions are made at the individual cell level and not in isolation, but rather in the context of a diverse community of interacting cells^2, 4, 5^.

Understanding biological “noise” and the potential functional role it plays in complex nonlinear processes, can explain the current spark of interest in single-cell analysis methods. In developmental biology, single-cell sequencing has been used extensively to investigate among others: relationships between different stem cell stages^6^, changes of the transcriptome from oocyte to morula in human and mouse embryos^7^, relationships of cell fate to gene expression from zygote to blastocyst^8^, characterization of stem cells in early embryos^9^. Likewise, in cancer research single-cell analysis methods have offered important insights into cell decision making^10^ in the context of a tumor’s microenvironment. Tumor heterogeneity results in cell lineages with vastly different molecular and histopathological phenotypes which regulate the aggressiveness, invasiveness, metastatic potential and resistance of the tumor to radiation and chemotherapy^11^. It is possible that cells heterogeneity contributes to the tumor’s adaptation to treatment leading to therapeutic failure^12^, so the capability to identify cell subpopulations becomes critical, since targeting the right one(s) might eventually prevent tumor’s growth.

Despite their differences, most single-cell data analysis methods are applied under the presumption that the population of cells analyzed is more or less homogeneous. However, this is far from being the case, since in reality cells of different phenotypes are mixed even in clonal cell populations. Methods to unmix subpopulations *in-silico* and infer their regulatory mechanisms have only recently started to emerge^13-20^. In Trott et al.^13^ the authors observed widespread heterogeneities in the expression of eight genes. Using correlation analysis they revealed the existence of three distinct stem cell states, distinguished based on the expression of Nanog (a pluripotency marker) and Fgf5 (a differentiation marker). Each state is associated to a collection of active sub-networks with differing degrees of node connectivity, correlated with self-renewal, primed-for-differentiation, and transition states respectively. Other studies have introduced the notion of a “trajectory”, i.e. a pseudo-time ordering of cells involved in a biological progression. For example, Trapnell et al.^14^ proposed an unsupervised learning algorithm (called Monocle) increasing the temporal resolution of transcriptome dynamics for RNA-Seq data collected at multiple time-points. In essence, cells are ordered in a way that maximizes the transcriptional similarity of successive cell pairs and a trajectory is formed by selecting the longest path of the constructed minimum spanning tree. In order to surpass limitations of Monocle, Haghverdi et. al.^15^ proposed a diffusion maps based approach for dimensionality reduction and cells ordering along the differentiation path. Another heuristic method (called SCUBA) introduced by Marco et.al.^16^, uses k-means clustering of the expression data, along with gap statistics to determine the number of classes (bifurcations) on a constructed binary tree of cellular hierarchy. It involves gene dynamics modelling for multi-lineage transition detection and assessment of related changes in gene expression patterns. In another work, Buettner et al.^17^ describe a computational approach using single-cell latent variable models (scLVM) to reconstruct the hidden factors from the observed data. Their model was used to assess the gene expression variance explained separately by the biological, technical, and hidden factors, aiming to reveal hidden subpopulations. In addition, Bendall et al.^18^ developed the so called Wanderlust algorithm, a graph-based approach ordering cells along a trajectory representing their most likely placement along a developmental continuum. Wanderlust determines a cell’s position based on edges (on the graph) between neighboring cells, where longer paths are less reliable than shorter ones. Moreover, Ocone et al.^19^ presented a framework that recovers the temporal behavior of cells using the Wanderlust algorithm^18^ and reverse engineers gene regulatory networks. Most recently, Setty et. al.^20^ introduced the Wishbone method which in essence is the evolution of the Wanderlust^18^. Cells are ordered using a graph-based approach based on distances that correspond to developmental chronology between cells. Briefly the workflow is: data ordering along pseudo-time, bifurcation identification, and cell to state association via proper marker selection. Overall single-cell gene expression profiling methods allow the identification of cell states and the reconstruction of networks involved in a biological procession (e.g. cell differentiation, tumor development etc.). As such, they have provided a blueprint for studying cell paths of evolvement and analyzing the behavior of reconstructed gene regulatory networks. Identification of cell subpopulations is a critical step in single-cell studies since it reveals information concerning the "logic" of dynamic state transitions, e.g. from pluripotency to differentiation, or from normal to diseased and to treated states. However, in all related works in trajectory definition or subpopulation identification^13, 14, 17, 18^ it is explicitly or implicitly assumed that cells progress smoothly and continuously in terms of transcriptional activity during state transitions.

We introduce here a novel computational methodology that does not make such an assumption, that is questionable as also noted recently in Moris et. al.^21^, but rather analyzes cell state transitions in a solid probabilistic framework, whereby expression signals modulate the probabilities of transition events. Specifically, cell subpopulations determination and trajectories construction take place in posterior probabilities space and not in genes expression space. In addition, our methodology is totally unbiased, relying solely on unsupervised machine learning methods, not assuming any knowledge regarding the identity and interactions of "key-player" genes governing each state transition. One can think of our approach as applying first *in-silico* cell-sorting to uncover first the major cell states (phenotypes) co-existing and interacting in a heterogeneous cell population, and then zooming-in and analyzing in detail each and every transition of interest among pairs of states, without relying on any cell type-specific gene markers.

For any A-to-B cell state transition that the user wants to study in detail, our methodology applies the following pipeline of computational steps: extracts from the dataset the subpopulation of cells most specific to the analysis of this transition; this is important since many state transitions may be “active” simultaneously in a dataset where several mixed cell subpopulations are interacting. Then, for the A-to-B state transition of interest, it constructs in probabilities space an ordered sequence (trajectory) of the cells involved in the biological progression of the transition. More than that, it partitions the extracted trajectory into three consecutive phases, called *micro-states* μstates). The ordered cells of the first μstate capture the process of departing from the source state-A, those of the second μstate represent the more dynamic transitory phase, and the cells of the third μstate represent the process of arriving to the destination state-B. In essence, we isolate from the heterogeneous dataset and place into a progression order the most relevant to the transition subpopulation, having only cells that show a preference to "move" from state-A towards state-B in the complex epigenetic landscape represented by the whole dataset. To make an analogy, it is like constructing a "staircase" in probability space having three regions μstates); the top padding (ground-A μstate), the stairs area (transition region μstate), and the bottom landing (ground-B μstate). Once the "staircase"-like trajectory is determined for the A-to-B transition of interest, our pipeline can: a) identify, in a parsimonious manner, the "key-player" (regulator) genes governing the transition, b) reverse engineer (infer) causal gene regulatory networks (GRNs) of the key-players for each microstate of the trajectory. The reverse engineered GRNs at the μstate level can provide information on how the interactions of key-players evolves during the transition progresses, thus providing useful insights (hypotheses) into the "logic" of the time-varying dynamical behavior of the biological system.

To the best of our knowledge this is the first methodology reported in the literature that, starting from the normalized gene expression profiles of a heterogeneous population of single-cells and for every state transition of interest, it can perform the following three important tasks: 1) Construct in probability space a trajectory specific to the state transition based on principled probabilistic methods not involving heuristic graph-based methods that process directly the noisy expression data, 2) Extract cell subpopulations μstates) representing each phase of the state transition under study, and 3) Identify among all genes in the dataset the most important regulators governing the specific transition and infer parsimonious regulatory networks of their causal interactions during each phase of the transition, all in an unsupervised, unbiased and data-driven manner. The same analysis can be repeated for all transitions, thus producing a probabilistic representation of the "epigenetic landscape" of the biological process captured by the available dataset.

## RESULTS

### The Computational Workflow

Figure 1 provides an overview of the proposed single-cell data analysis workflow. In summary, we first apply dimensionality reduction using Principal Component Analysis^22^ (Fig. 1b). Then Gaussian Mixture Modeling^23^ (GMM) is applied (Fig. 1c) to (i) identify the most likely number of cell states (major cell phenotypes) mixed in the dataset, (ii) detect outlier cells to be excluded from downstream analysis, and (iii) determine the posterior probability distribution for each cell to "belong" to each one of the identified cell states (major phenotypic categories) as suggested by the data. We remark that the ground truth (cell type) information used to color cells in Fig. 1b is used only for visualization purposes and it is not utilized at all in the totally unsupervised GMM analysis. The GMM analysis actually suggests the existence of 4 cell states, i.e. one for each cell type (HSC, LMPP, PremegE) mixed in the analyzed dataset^24^ (see Online Methods) plus a fourth class encapsulating outlier cells (see Supplementary Material file, Figure SM1) that are excluded from any downstream analysis. Let’s assume, without loss of generality, that we now want to focus on the analysis of a specific state transition, say from a source state-A to a destination state-B. We first construct a trajectory in posterior probabilities space including only cells that are relevant to the study of this specific state transition (Fig. 1d). Moreover, we partition this trajectory into three successive microstates μstates), i.e. three cell subpopulations, called "ground-A", "transition", and "ground-B" (details in Online Methods), representing the three phases of a state transition progression. Then, we analyze the gene expression profiles of the trajectory cells to identify the genes that are the most "active”, or so called "key-player" genes (Fig. 1e) in terms of driving the state transition (algorithmic details in Online Methods). Finally, we employ the well-known GENIE3 algorithm^25^ to infer μstate level gene regulatory networks of key-player genes, and use correlation analysis to determine the sign (excitatory vs. inhibitory) of the inferred causal network arcs (Fig. 1f).

**Figure 1.**
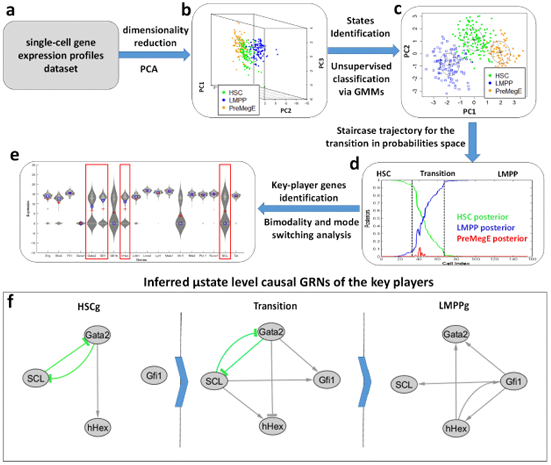
Overview of the Single-cell Data Analysis Workflow using the analysis of the HSC-to-LMPP state transition as an example. a) We start with a heterogeneous single-cell gene expression profiles dataset. b) Apply dimensionality reduction using Principal Components Analysis. c) Apply unsupervised clustering to determine major cell states and the posterior probabilities of cells to belong to each state using Gaussian Mixture Modeling and model selection. d) Select a specific state-A to state-B transition of interest to analyze (HSC-to-LMPP in our example without loss of generality). Construct the corresponding "staircase" trajectory of cells in posterior probabilities space, "descending" from the source to the destination state. In addition, define a partition of this trajectory into three consecutive regions (and corresponding ordered cell subpopulations), called *microstates* μstates); namely HSC- ground (HSCg), Transition, and LMPP-ground (LMPPg) μstates in our case. e) Identify the "key-player" genes governing the dynamics of the state transition by applying bimodality and expression mode switching analysis. f) For each μstate reverse engineer causal gene regulatory networks of the key- player genes; only the top-scored interactions are shown (see text and Online Methods for details). The same procedure can be repeated for any transition of interest defined by a pair of identified states.

### Application to hematopoietic dataset and the analysis of the HSC-to-LMPP transition

Moignard et al.^24^ analyzed the expression of 18 densely interconnected transcription factors in a population of 597 single-cells belonging to 5 primary hematopoietic stem and progenitor cell subpopulations. They clustered the cells according to expression profiles and performed correlation analysis to infer a gene regulatory network. The selected transcription factors revealed not only characteristic expression states for the different cell populations, but also previously unrecognized regulatory relationships^24^. This included a putative regulatory triad, consisting of the Gata2, Gfi1 and Gfi1b genes, which was validated using transcription and transgenic mouse assays. We demonstrate the proposed methodology using this publicly available dataset^24^, focusing in particular on the analysis of the important HSC-to-LMPP state transition. To this end, we selected 350 cells, those with HSC, PreMegE and LMPP cell types (determined by cell-sorting), which comprise the root and its two children in the top-level of the hematopoietic system cells hierarchy, as presented in Moignard et. al.^24^.

By applying dimensionality reduction using PCA (Fig. 2a) we found that the first three principal components (PCs) can explain ~39% of the total data variance in the dataset (Fig. 2b). It is interesting to observe that LMPP and PreMegE cells fall on opposite sides with respect to the HSC cells (especially along PC2) something that complies with our intuition (Fig. 2d). Next, the reduced 3-dimensional (3D) data vectors are clustered (in PCA space) using unsupervised Gaussian Mixtures Modeling (GMM)^23^. The GMM model exhibiting the best performance has 4 components, one representative of each actual cell type and an additional one representing outlier cells (see black points in Supplementary Information, Figure SM1). We exclude outlier class cells from the analysis and re-compute the posteriors for the rest of the cells (Fig. 2c, details in Online Methods). In Figure 2e we provide the "confusion matrix" that can be constructed since the original cell types (labels) are known for this dataset. We observe that PreMegE and LMPP type cells are not confused for each other. This indicates that the clustering was effective, since it is possible to have some HSC type cells resemble to LMPP or PreMegE cells and also some PreMegE or LMPP type cells (not “mature” enough) resemble to HSC cells, but the PreMegE and LMPP cell fates are quite "orthogonal" to each other with very distinct transcriptional phenotypes. These results indicate that even if some cells are known to belong to a certain cell type (after FACS sorting) they may have already started progressing towards another state or their expression may be an outlier with respect to that state. A data-driven methodology aiming to reconstruct GRNs that capture the dynamics of a specific state transition should take into account the presence of inherent biological and technical noise in single-cell expression profiles datasets. GMM based clustering meets this requirement in a principled manner, since it recovers the full posterior distribution for each cell to belong to each one of the classes suggested by the data in an unbiased manner (soft computing). In addition, it can be used to identify outlier cells that are not represented clearly by any of the major cellular phenotypes as suggested by the data.

**Figure 2.**
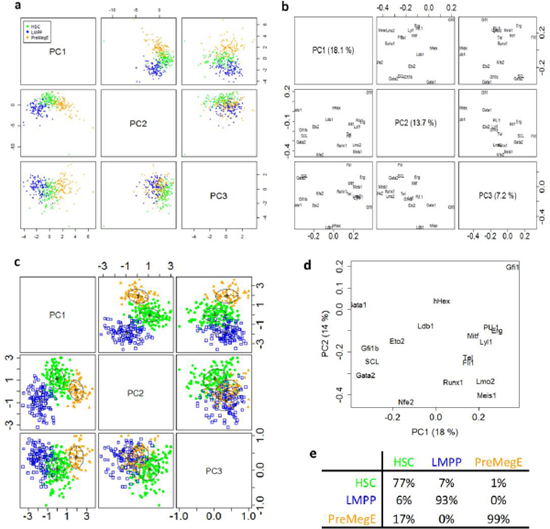
Dimensionality reduction followed by unsupervised Clustering. a) Principal Components Analysis of single-cell gene expression profiles in a cell population formed by mixing three cell lineages (cell-types denoted by color). b) PCA loadings plot showing the impact of individual genes (variables) to the distribution of cells (samples). c) The result of unsupervised clustering in PCA space using Gaussian Mixture Modeling and best model selection; three classes are identified which actually matches the number of true cell lineages. d) Loadings plot for PC1 and PC2 showing the impact of individual genes to the distribution of cells. e) Confusion matrix showing the percentages of true cell types (columns) for each identified (computed) cell class (row). The true cell type labels are not used in any step of the totally unsupervised machine learning methodology.

Let’s now focus without loss of generality on the analysis of the HSC-to-LMPP state transition of interest and identify the population of cells forming a staircase-like trajectory in probability space, leaving HSC with a direction towards LMPP (see Online Methods) and its three μstates. This actually results in producing a trajectory and its three-subset partition consisting of: the ground-HSC (HSCg) μstate (33 cells), the Transition μstate (34 cells) and the ground-LMPP (LMPPg) μstate (88 cells). The ordering of the HSC-to-LMPP staircase trajectory cells based on their posterior probabilities and the μstate boundaries are shown in Figure 3. Using only these ordered trajectory cells, we determine next the “key-player" genes, i.e. those genes that not only exhibit clear bimodal expression but also switch modes of expression as the trajectory crosses from the starting (HSCg) to the ending (LMPPg) ground μstate (see Online Methods). Using this analysis, the key-player genes turn out to be the tetrad Gata2, Gfi1, hHex, and SCL (also known as Tal1) as shown in Figure 4a. Their expression patterns along the trajectory are shown in Figure 4b-e. Interestingly, the first two transcription factors (Gata2 and Gfi1) are also identified as key genes in Moignard et. al.^24^. Gfi1 is a transcription repressor essential for hematopoiesis^26^. SCL

**Figure 3.**
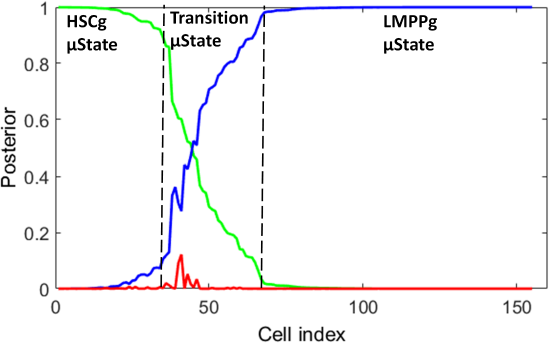
Posterior probabilities for the subpopulation of cells most representative of the HSC-to-LMPP state transition to be analyzed. The cells are ordered on a "staircase"-like trajectory in probability space in decreasing posterior probability to the source state (HSC, green curve). For the ordered cells, the blue curve shows their posterior to the destination state (LMPP). The third curve (red) shows the posterior probability of the trajectory cells to the third state (PreMegE); by construction of the trajectory it has the smallest value for every cell in the subpopulation since this state is neither the source not the destination state for the transition under study. Dashed vertical lines mark the three microstate ( μstate) boundaries. (See text and Online Methods for details).

**Figure 4.**
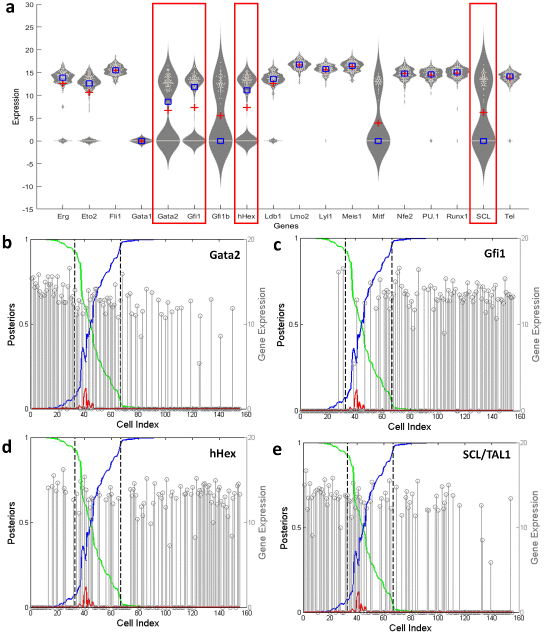
"Key-player" genes identified for the HSC-to-LMPP state transition. a) Violin plots of all measured gene expression profiles. Genes with violin plots in rectangular boxes are the key-players selected using bimodality and mode switching analysis criteria (see Online Methods for details). b-e) Expression patterns of the four key-player genes Gata2, Gfi1, hHex, SCL for the ordered cells of the "staircase" trajectory. As we can see, Gata2 and SCL are "firing" mostly in the source HSCg μstate, while Gfi1 and hHex in the destination LMPPg μstate (see text and Online Methods for details).

(TAL1) is required for the generation of all hematopoietic lineages and is essential for establishing the transcriptional program responsible for the formation of hematopoietic stem cells^27^. The homeobox gene hHex is a very important gene which has been found to regulate the earliest stages of definitive hematopoiesis^28^ as well as the proliferation and differentiation of hemangioblasts and endothelial cells during ES cell differentiation^29^. In addition, hHex was found not to be required for HSC function in adult hematopoiesis, but to play a critical role in early lymphoid specification^30^.

We remark that by inspecting the PCA loading plots of Figure 2b we can establish a visual impression into the “importance” of these genes to the GMM clustering result and therefore to the downstream analysis. At the plane defined by the first two principal components (Fig. 2d), and Gfi1 roughly lie at the opposite ends of the main diagonal. We also observe that if we start from HSC cells (green in Fig. 2a top middle panel) and move in a direction parallel to the diagonal we arrive at LMPP state cells (blue in Fig. 2a). So, it is intuitively expected that these genes will be important for this state transition. We should emphasize that unlike other single-cell expression data analysis methodologies, all outcomes here have been produced in a totally unbiased and unsupervised manner.

In the last step, we use a well-established tool, GENIE3^25^, to infer causal gene regulatory networks for each μstate, as shown in Figure 5. Only the top-scored interactions are shown in the Figure (see Online Methods). Colors are used to mark gene interactions according to whether they are top-scored in one or more μstates of the transition. Being able to zoom in and infer μstate level GRNs, allows to shed light on the possible role of key-player genes during the state transition process. The top-scored interactions may change dynamically as the HSC-to-LMPP transition process progresses. In particular, from the inferred GRNs we can observe that in the HSCg μstate there is a regulatory module of two key-player genes, {GATA2, SCL}, with bilateral negative interactions between them, and a positive interaction Gata2-to- hHex, suggesting an increasing role for hHex as cells start transitioning towards the LMPP state. During the transition μstate, the SCL-to-Gata2 loop persists; in addition, new regulatory links emerge connecting both its members to the other two genes, hHex and Gfi1 (all positive except Gata2-to-hHex). This can be thought as the emergence of a major role for Gfi1 and hHex as cells move towards the LMPP state, initiated by the other partners, Gata2 and SCL, as cells depart from the HSC state. At the terminal LMPPg μstate of the transition, a new loop {Gfi1, hHex} emerges, with one of its members, Gfi1, interacting with all other genes. It is also interesting to observe that the direction of the interaction of hHex and Gata2 is reversed relatively to the beginning of the transition in a manner resembling a request-acknowledge “handshaking” communication protocol in digital circuits^31^. It is evident that the hierarchical analysis strategy we have applied (from the whole dataset to major states, to trajectories connecting states, to μstates of individual trajectories) allows us to zoom in and bring on the surface dynamic sub-networks that not only reveal the key-players for any state transition of interest in a parsimonious manner, but also help us reconstruct possible scenarios (make interesting hypotheses) suggested by the data as to the dynamic evolution and timing of predominant interactions. Our parsimonious modeling enables analyzing all phases of a transition at any desirable level of resolution while remaining focused on identifying the main interactions among only the most relevant to the transition gene players.

**Figure 5.**
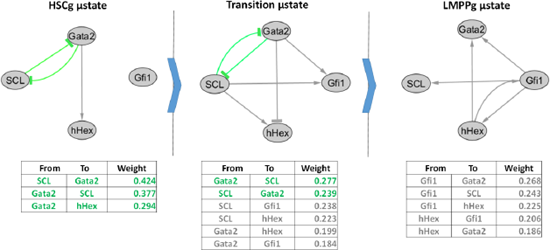
Causal Gene Regulatory Networks inferred for the three μstates of the HSC-to-LMPP transition using the GENIE3 algorithm. From left to right: HSCg, Transition, and LMPPg μstate networks having as nodes the four key-player genes (see Figure 4). Only the top-scored interactions are shown for each network in descending order of GENIE3 weights (see text for details). Green interactions appear in HSCg and persist in Transition μstate; gray interactions are present only in the top-scored list of one μstate network.

Our findings are validated by the related literature. Wilson et al.^32^ indicated that SCL (Tal1) and Gata2 regulate Gfi1 expression. Here we see that both Gata2 and SCL affect Gfi1 in the transition μstate, where no apparent influence of Gfi1 from SCL or Gata2 exists in the HSCg μstate. However, the dyad {Gata2, SCL} seems to be self-regulated in such a way that in the transition μstate both Gata2 and SCL affect the expression of Gfi1. In published ChIPseq data of HSC7 cells (representative of stem/progenitor cells) available at the ChIPseq repository Codex [http://codex.stemcells.cam.ac.uk], there is a peak at the SCL promoter and a peak at the Gata2 -83kb enhancer. Likewise, there is an SCL related ChIPseq evidence that shows binding at the Gata2 locus (including -83kb enhancer) and Gfi1 locus (-35kb enhancer in the neighboring Evi5 gene) identified in Wilson et al.^32^. Moreover, Pimanda et al.^33^ also provided evidence of interaction between Gata2 and SCL, which along with Fli1 form a triad regulating the development and differentiation of hematopoietic cells. It is also known that GATA2 is required for HSC survival and self-renewal, interacting with a complex network of transcription factors that specify early lineage commitment, including SPI1 (PU.1), FLI1, SCL (TAL1), LMO2 and RUNX1, among others^34-36^. Moreover it has been shown that Gfi1 expression is progressively upregulated in MPP1, MPP2, and LMPPs progenitors^37^. In addition, Gfi1 is expressed in lymphoid precursor cells such as CLPs and ETPs that settle the thymus and, subsequently, in the early stages of T- and B-cell development^38-41^ but also in GMPs and in monocytes and granulocytes^37^. In Schütte et al.^42^, the authors generated a network using the Biotapestry^43^ software, where all the aforementioned interactions are confirmed, in terms of corresponding binding sites; i.e. the regulatory dyad {Gata2, SCL} exists and seems to play an important role in hematopoiesis. Finally, in Jackson et al.^30^ a stage-specific role for hHex during hemangioblast development in embryogenesis was identified, placing hHex in the class of transcriptional regulators that have roles in commitment and/or expansion of HSCs during embryogenesis, but are dispensable for HSC maintenance in the adult hematopoiesis, including SCL/Tal1^44^ and Runx1^45^.

### Reconstructing a probabilistic view of the epigenetic landscape

In addition to studying in detail a specific transition, the proposed methodology can be used to reconstruct a probability-based representation of the “epigenetic landscape” of state transitions represented by a mixed single-cell population. The dataset used to demonstrate this analysis is the whole hematopoietic dataset from Moignard et al.^24^ consisting of 597 single-cell expression profiles belonging to 5 primary hematopoietic stem and progenitor cell types; namely HSC, PreMegE, LMPP, CLP and GMP. Considering the dataset as unlabeled (i.e. by not exploiting in any way the available knowledge regarding cell type labels) and following the presented analysis workflow, all major states (subpopulations in Moignard et al.^24^) were identified (refer to Figure SM2) and all state-to-state transition trajectories were extracted (see Online Methods). In summary, after applying dimensionality reduction where only the first 3 principal components are taken into account, unsupervised GMM model selection identifies a model with 5 classes (states) as being the “best”, matching the number of the true underlying cell subpopulations (refer to Figure SM2). Then, all possible trajectories, starting from a state-*A* and going to any other state-*x* (for all pairs of state transitions) were extracted and a transition probability *P(A →x)* is calculated as the ratio of the number of cells *c(A,x)* having highest posterior probability to the origin state-A and also belonging to the *A-to-x* trajectory, to the total number of cells *c(A)* with highest posterior to state-*A*. Intuitively, this ratio represents an estimate of the conditional probability that cells originating in state-*A* will transition to state*-x* and can be used to generate a network view of the transitions represented in the whole dataset. Figure 6 shows this view of the "epigenetic landscape" where the edge weights have been normalized to sum up to 1 by scaling their original values (i.e. dividing by 5, the total number of states). Each edge weight A-B is computed as the sum of the two transition probabilities *P(A →B)* and *P(B→A).* As we can notice, the most likely transitions supported by the analysis are HSC-PreMegE, HSC-LMPP, LMPP-CLP and LMPP-GMP. These are indeed the transitions recognized as major throughout the literature and in Moignard et al.^24^. The rest of the transitions exhibit much lower probability values: Some are practically zero (i.e. < 1%, e.g. PreMegE-GMP and PreMegE-LMPP) and are not shown in Fig. 6, while some others are small-valued, e.g. PreMegE-CLP and CLP-GMP. For the former (practically zero probability) case, the transitions from PreMegE (which is a megakaryocyte-erythroid lineage) towards either GMP (myeloid lineage) or LMPP (lymphoid lineage) are highly unlikely to occur. For the latter case, we could think that the initiating/early cells in these trajectories are at a latent/intermediate μstate that would “allow” the corresponding transition. For example, for the CLP-GMP transition the cells with highest posterior either in CLP or GMP state could actually be in LMPP state (“mother” state for both), allowing the transition to happen. Correspondingly for the PreMegE–CLP transition trajectory, some of the cells may actually belong to a latent MPP state which is prior to PreMegE differentiated state. Concerning the low probability transitions we should note that as discussed in Athanasiades et al.^46^ hematopoiesis exhibits unsynchronized development, where each cell may demonstrate a different degree of differentiation along the differentiation continuum. Therefore, in principle no transition is “prohibited” (zero probability), but rather a probability assignment to all transitions is a more appropriate (physiologic) treatment capturing the inherently stochastic nature of biological processes.

**Figure 6.**
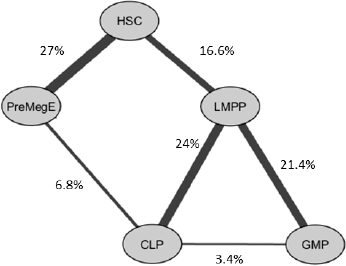
Network representation of the Epigenetic Landscape extracted based on the analysis of the whole hematopoietic dataset: Each undirected edge A-B models two directed edges A→B and B→A with the edge weight corresponding to the sum of the two state transition probabilities. A state transition probability is estimated as the percent of cells of the originating state that are also on the trajectory towards the destination state (see Online Methods). Edge weights have been normalized to sum up to one for all transitions in the network and weights <1% are not shown in the Figure. The higher the edge weight the thicker the corresponding network edge line. The well-known 3-level binary tree structure of the hematopoietic hierarchy of cell lineages is revealed by the unsupervised data analysis (thick line edges). Some other relatively small probability transitions are also possible (see text for details).

## DISCUSSION

We have presented an end-to-end computational methodology for the analysis of single-cell gene expression datasets that for every state transition that the user wants to analyze it can: 1) identify the subset of cells (sub-population) in the dataset most relevant to the analysis of the selected state transition, 2) order these cells along a staircase-like trajectory as they transition in posterior probabilities space, leaving the source and going towards the destination state, 3) identify in a parsimonious manner the key-player genes interacting actively during the state transition, and 4) infer actionable causal GRNs of these genes, capturing also the time varying network characteristics at the three phases μstates) of a state transition; departure from the source state, mid-way in the transition, arrival to the destination state. All the above analysis is performed in a totally unbiased, data-driven, unsupervised manner and is fully automated.

We have presented the methodology through the analysis of the fundamental HSC-to-LMPP state transition using a publicly available single-cell gene expression profiles dataset. We obtained results that are in accordance with the literature while also offering new insights on the key-player genes and how their interactions may evolve dynamically during the transition. The presented methodology does not depend on the process that generates the data and can be applied to single-cell expression profiles produced by either fixed-time or time-course experiments. Moreover, the concept of cell state as used here can be interpreted very broadly. For example, in a dataset with untreated and treated tissue cells, source state-A can be the state prior to treatment and destination state-B the state after the treatment; or state-A can be healthy cells while state-B tumor cells in the same tissue, etc.

Recently, the exploration of micro dynamics in cancer has emerged as a very important topic^47, 48^. There is a tremendous interest in dissecting the micro-heterogeneity inherent in tumor cell populations that exhibit a wide phenotypic variability in early cancer stages. It has long been appreciated that tumors are composed of heterogeneous cell populations, with a pool of cells that display stem cell properties and drive the evolution of a tumor to a gradually more aggressive phenotype^47, 48^. Recent studies have shown that circulating tumor cells (CTCs) isolated from breast, colon, and hepatic cancer patient blood are similarly composed of a heterogeneous pool of cells, some of which have tumor-initiating/stem cell properties and display epithelial–mesenchymal transition features and low apoptotic propensity^49-51^. For scientific and clinical purposes, it is very important to understand the dynamics of heterogeneous tumor cell populations and characterize their variability across different phenotypes, specifically under radio- and/or chemotherapy^11^ conditions. Moreover, it is of fundamental importance to understand the molecular basis of heterogeneity. So far, heterogeneity in cancer cells is approached by means of finding appropriate markers, a process which apart from being expensive and time consuming is also inefficient since in early tumor stages cancer cells are very well "camouflaged" and the specific markers may not be expressed^11^. As discussed in Freeman et al.^52^, marker based determination of subpopulations is not always applicable and a marker-free approach for CTCs isolation and subpopulation definition would be very beneficial. We believe that our approach provides a general data analysis framework that can be exploited in this direction to recognize cancer cells subpopulations and characterize their properties, such as significant genes/transcription factors interacting, parsimonious GRNs, identification of top-scored interactions and how they evolve as the subpopulation mix is changing.

Recently, Moris et al.^21^ have suggested an alternative view of Waddington’s epigenetic landscape^53^ challenging the notion that transcriptional expression follows a smooth/continuous path as cells make state transitions, a hypothesis that underlies many current pseudo-temporal trajectory construction methods. According to Moris et al.^21^, cells are rather following a stochastic pattern of gene expression where transcriptional signals modulate the probability of state transitions. In line with this more plausible view, our methodology does not assume expression smoothness during the transition and takes into account the inherent stochasticity in the underlying biological mechanisms since our state transition trajectories are "crafted" in the space of posterior probabilities to states. To the best of our knowledge, our methodology is unique among all other related methods (regarding pseudo-time organization of cells) in that it can reconstruct a probabilistic representation of the epigenetic landscape of state transitions involved in the underlying biological process as suggested by the available dataset.

Finally, we should emphasize that although our methodology was presented here using an RT-qPCR single-cell dataset (commonly used throughout the trajectory related literature), this does not present a limitation, since the same approach can be used to analyze normalized single-cell RNA-Seq gene expression profile datasets. What we expect in such a case is that we may identify a larger number of key-player genes. This may require an additional pre-filtering step to appropriately rank-order and reduce the number of key-player genes down to a reasonable number before inferring GRNs, especially if the dataset is not having a sufficiently large number of cells.

## ACKNOWLEDGMENTS

We would like to thank Dr. Vicki Moignard and Prof. Bertie Gottgens, University of Cambridge, Cambridge Institute for Medical Research, UK, for providing feedback regarding inquiries on the dataset used and the results.

## METHODS

The proposed single-cell data analysis workflow is summarized in Figure 1. Let’s assume that we are given a dataset of normalized gene expression profiles for a heterogeneous population of single-cells. Each cell is represented in the dataset by a vector of gene expressions, with the vector dimensions matching the number of genes. Our methodology does not assume any prior knowledge of the individual cell types (lineages), on the contrary it treats the population as a heterogeneous mix of cells of unknown origins. First, we seek to cluster together cells exhibiting similar expression profiles (i.e. belonging to the same cell state) irrespectively of their lineage of origin and also identify outlier cells with expression profiles that are drastically different from all other cells. However, before applying unsupervised clustering of cell expression profiles, we perform first dimensionality reduction to compensate for possibly a small ratio of samples (cells) to variables (genes measured) in the dataset.

### Dimensionality Reduction and Unsupervised Clustering to identify the major cell states in the population

Principal Components Analysis (PCA)^23^ (using the *princomp* function in R [https://stat.ethz.ch/R-manual/R-devel/library/stats/html/princomp.html]) is applied first for dimensionality reduction. The PCA results (first three principal components) for the dataset including the cells of 3 cell types, HSC, LMPP and PreMegE, are provided in Figure 2a and the corresponding PCA loading plots are shown in Figures 2b and d. PCA results for the whole dataset used in Moignard et al.^24^ (i.e. including also the cells with cell types CLP and GMP) can be found in Supplementary Material Figure SM2. Gaussian Mixture Modeling^23^ (GMM) using the EM algorithm^54^ is applied next, complemented with best model selection, as implemented by the *mclust* package in R [https://cran.r-project.org/web/packages/mclust/mclust.pdf]. For the Gaussian components of the mixture we assume that they are spherical in shape (i.e. their covariance matrices are diagonal) with unequal volumes. The former assumption is justified because in lack of any other information to the opposite, spherical Gaussians are expected to consolidate cells into subpopulations that do not intrude into neighboring subpopulations. The latter assumption is intuitively justified since in general we cannot assume that the mixed subpopulations are of equal size. Figure 2c (and Figure SM2e,f) shows the best model selection using the Bayesian Information Criterion^55^ (BIC). We consider as optimal, the model with the smallest number of components (minimal complexity) that exhibits a near maximum BIC. This compromise is a way of selecting the best model in a parsimonious manner (see Figure SM3). Although every sample (cell) has a posterior probability to every identified class (soft computing approach), in Figure 2c (and Figures SM2e,f) it is colored based on the dominant class, i.e. the one with the largest posterior probability. We observe that the best model: (i) identifies as many classes as the number of true underlying cell types that were mixed to produce the dataset, and (ii) can detect outliers when they are apparent (see Figure SM1). A class is considered to representing outliers if it has a small number of cells in comparison to the total number, and its sample variance is comparable to the variance of all samples in the dataset. If an outlier class is identified by GMM, all its members are removed from the dataset and the posterior probabilities are then recalculated using the remaining cells (Figure 2c). Figure 2e provides the resulting "confusion matrix" that validates the clustering results. We observe that more than 77% of the green cell are actually HSC type cells. Since HSC type cells may evolve to assume either an LMPP or a PreMegE phenotype it is expected that in the green extracted class we may also find some “immature” cells from these lineages. In the case of unsupervised clustering of the whole dataset (all 5 cell types included) there was no outlier class detected. A probabilistic soft-computing framework for clustering samples, as the one employed here, is very appropriate to model the situation where a cell in a heterogeneous population may be in transition, and not clearly belonging to any specific phenotype but rather having a "membership" to different phenotypes. This modeling approach is fully exploited by our methodology when it comes to constructing a "staircase-like" trajectory of cells undergoing a state transition in posterior probabilities space.

### "Staircase" trajectory formation and definition of microstates

The focus of this work is on the reverse engineering of gene regulatory networks of key-player genes that are specific to and can inform a state transition of interest. For this purpose, we exploit the estimated posterior distribution, i.e. "how much” each cell belongs to each state as determined by the best fit GMM model, given the data (gene expression profiles). Since we focus on the analysis of one transition at a time, say from state-A to state-B, we exclude from the A-to-B trajectory cells with a significant posterior to any other third class *i* (different than A or B) satisfying:

*Prob(class_i_|data) > Prob (class_A_|data),* OR *Prob(class_i_|data) > Prob (class_B_|data).*

We then order the remaining cells (i.e. the subpopulation most relevant to the A-to-B state transition) in descending order of posterior probabilities to the source state-A. In this way we are effectively constructing a "staircase-like" trajectory of cells "descending" from (i.e. leaving) state-A, and going towards state-B in probability space, as depicted in Figure 3.

In Figure 3 we notice that the "staircase-like" trajectory is partitioned into distinct regions (cell subsets); the early one with cells exhibiting high posterior probability to the source state-A (high values for the green curve), the late one with cells exhibiting high posterior probability to the destination state-B (high values for the blue curve), and one in between with cells having considerable posteriors to both class-A and class-B. In order to determine in an unbiased manner the boundaries best delineate these three regions, to be called *microstates* μstates), we apply an algorithm that relies solely on the computed posterior probabilities. Specifically, on the curve of the class-A posterior (green) we determine two "knee-points" (high and low knee) marking the μstate boundaries. Each knee-point is the intersection of two lines that are obtained as a result of an iterative curve fitting procedure. Specifically, by "walking" along the curve (a fitted smoothing spline^53^ to the posteriors data) from left to right one bisection point at a time and fitting two lines, one to all points to the left of the bisection point and another to all points to the right of the bisection point, the left knee is the bisection point for which the sum of the Root Mean Square Errors (RMSE) for the two lines is minimized. This procedure is repeated, but now "walking" along the same curve on the opposite direction, from right to left, in order to define the second (bottom) knee-point. The resulting two knee-points Estate boundaries) are shown in Figure 3 using vertical dashed lines.

### Key-player genes identification for the state transition of interest

Once the trajectory of cells and the three μstates (called ground-A, transition, ground-B) have been defined for the A-to-B state transition of interest, we can identify the genes that exhibit “interesting” behavior along that trajectory and can be considered as the "key-players" with respect to the corresponding state transition. Let us consider the distributions of all measured gene expressions for the subpopulation of cells on the A-to-B trajectory, depicted using "violin plots" in Figure 4a. We consider a gene x to be a "key-player” gene for the A-to-B transition if the following two conditions are both satisfied:

1. Gene x expression profile exhibits a clear bimodal form, 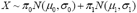, and it holds that 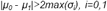 *μ_i_, σ_i_* are the mean and standard deviation of the two mode components, π_i_ are the mixing proportions, and *N()* is the Gaussian density function.
2. A mode of expression is clearly dominant in the ground-A μstate or in the ground- B μstate. Let us consider gene expression for a cell to be "high level" (denoted as level *j=1)* if it holds that gene expression ≥ μ_1_ – 2σ_1_, and "low level" *(j=0)* if gene expression ≤ μ_0_ ± 2σ_0_. Let *n_ij_* be the number of trajectory cells in μstate *i* (where *i* = {ground-A, ground-B}) that have expression level *j* for gene *x.* Let 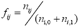 be an estimate of the probability that gene x exhibits expression level *j* at μstate *i.* Then we check if the following condition is satisfied: 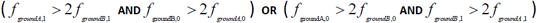 Intuitively, if for some gene *x* the percentage of cells exhibiting high (low) expression in the source μstate ground-A is greater than twice the corresponding percentage of cells in the destination microstate ground-B, and at the same time the percentage of cells exhibiting low (high) expression level at ground-B is twice the corresponding percentage of cells in the source microstate ground-A, we consider that gene x is making a high-to-low (low-to-high) expression transition along the trajectory i.e. switching modes of expression along the trajectory.

Genes that satisfy both conditions 1 and 2 above (i.e. have a bimodal expression profile and are switching mode of expression along the trajectory) are considered as "key-players" in the regulatory mechanism of the state transition, since their expression exhibits a distinct pattern between the two ground μstates (source and destination) showing that they play a major role during the transition. The key-player genes are provided in Figure 4 for the HSC-to-LMPP transition. Their expressions exhibit an interesting profile since it is not only bimodal (see Figure 4a: genes into red rectangles) but also switches expression mode along the trajectory, i.e. gene "firing" is a lot more frequent in one ground μstate than the other (see Figure 4 (b-e). It is clear that two of the identified key-player genes exhibit higher “firing” at the origin pstate (Gata2 and SCL) while the other two at the destination pstate (hHex and Gfi1).

### Gene Regulatory Networks Inference at the μstate level

At the last stage of the methodology we use the extracted trajectory information (cells belonging to each μstate) to infer gene regulatory networks (GRNs) for each μstate. To that end we employ the well-known GENIE3^25^ algorithm for networks inference. GENIE3 decomposes the inference of a regulatory network of *p* genes into *p* different nonlinear regression problems, where the expression pattern of each gene (target) is predicted from the expression patterns of all the remaining (p-1) genes using tree-based ensemble methods and in particular Random Forests^56^. The influence weight of a gene in the prediction of the target gene expression pattern is taken as an indication of a putative regulatory link. Putative regulatory links are then aggregated over all genes to provide a global ranking of the interactions from which the whole network is reconstructed. The algorithm was selected over other methods since it was the best performer in the DREAM4 [http://dreamchallenges.org/project/closed/dream4-in-silico-network-challenge/] and DREAM5 [http://dreamchallenges.org/project/closed/dream-5-network-inference-challenge/] in silico Multifactorial challenges. Moreover, GENIE3 does not make any assumptions on the nature of the regulation, can deal with complex non-linear interactions, and infers directed (causal) interactions, unlike correlation networks where none of the aforementioned desirable properties hold. Since GENIE3 produces weights of relative importance for all interactions, we applied a simple criterion for selecting the most important ones. Specifically, we retain only the top-scored network connections having weights larger than the maximum value of all network weights minus their standard deviation: *(max(weights)-std(weights)).* In addition, we perform correlation analysis at each μstate in order to infer the sign (activation or inhibition) of the inferred network link. The correlations are calculated in R, using the *cor.ci* function of the *psych* package [https://cran.r-project.org/web/packages/psych/psych.pdf]. We computed the spearman correlations with bootstrapped confidence interval and p-values<0.01. Figure 5 presents the GRNs inferred for the three μstates for the HSC-to-LMPP state transition.

### Reconstructing a probabilistic view of the epigenetic landscape

Finally, we applied our methodology to the whole dataset (all cell types included) and extracted trajectories for all possible combinations of source and destination states (state transitions). At this point we calculate the ratio of the number of cells that have highest posterior probability to the origin state-A and also belong to the trajectory leading to destination state-x *(c(A, x)*) to the total number of cells with highest posterior to state-A, *c*(*A*), i.e.:

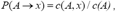

Where x can be any state other than A. This ratio provides an estimate of the transition probability *P(A →x)* to state-x of cells in state-A and can be used to generate a probabilities-based network representation of the "epigenetic landscape" as suggested by the dataset. By construction of the trajectories of transitions *A-to-x,* it is guaranteed that for any given source state-A the sum of transition probabilities *P(A →x),* over x any state other than *A,* is equal to 1. So *P(A →x)* provides an estimate of the conditional probability P(next state is x | current state is *A).*

#### Materials

The presented framework has been evaluated using the hematopoietic single-cell expression dataset from Moignard et al.^24^ (see Supplementary Table S3 at http://www.nature.com/ncb/journal/v15/n4/full/ncb2709.html). In this paper the authors performed single-cell gene expression analysis using 48.48 Dynamic Array integrated fluidics chips (M48, Fluidigm) on the BioMark HD platform (Fluidigm) over 24 genes for 597 cells in order to characterize the transcriptional networks in blood stem and progenitor cells. The selected gene set included 18 transcription factors, 5 housekeeping genes and the stem cell factor receptor *c-Kit.* The data set of 597 cells consisted of a mixture of the following cell types: HSCs (121 cells), pre-megakaryocyte/erythroid progenitors (PreMegE 113 cells), lymphoid primed multipotent progenitors (LMPP 116 cells), and granulocyte-macrophage progenitors (GMP 124 cells). Prior to the analysis we followed the data preprocessing steps presented in Moignard et al.^24^. Next, since the maximum data value of 15 was set by Moignard et al.^24^ to represent no expression, we inverted the expression values by subtracting them from the maximum value (i.e. 15), so that high (low) values correspond to high (low) levels of gene expression. For analyzing the HSC-to-LMPP state transition we used only the HSC, LMPP and PreMegE cells (350 in total) in Moignard et al.^24^. However, for reconstructing the probabilities-based network representation of the epigenetic landscape we used the whole dataset.

#### Data Availability

The dataset used in this study is from Moignard et al.^24^ and is available as part of its supplementary information file (Supplementary Table S3), see http://www.nature.com/ncb/journal/v15/n4/full/ncb2709.html.

